# Z-TAC enables custom and combinatorial degradation of cell surface proteins

**DOI:** 10.64898/2026.04.03.716357

**Authors:** Dingjingyu Zhou, Lawrence Shue, Sean Gao, Eric S. Fischer, Ryan A. Flynn, Xin Zhou

**Affiliations:** Department of Cancer Biology, Dana-Farber Cancer Institute, Boston, MA, USA; Department of Biological Chemistry and Molecular Pharmacology, Harvard Medical School, Boston, MA, USA; Stem Cell and Regenerative Biology Program, Division of Hematology Oncology, Boston Children’s Hospital, Boston, MA, USA; Department of Stem Cell and Regenerative Biology, Harvard University, Cambridge, MA, USA; Harvard Stem Cell Institute, Harvard University, Cambridge, MA, USA

**Keywords:** Targeted protein degradation, cell-surface proteins, antibody engineering, multi-pass proteins, transferrin receptor 1, endocytosis, combinatorial targeting, membrane protein signaling

## Abstract

Cell-surface degrader platforms typically require target-specific engineering and have therefore been applied to a relatively small set of protein targets. Here we report Z-TAC, a strategy that enables plug-and-play conversion of existing IgG antibodies into cell-surface protein degraders. Across multiple targets from distinct protein families, Z-TAC induced efficient and sustained degradation of both individual receptors and receptor combinations. For a multi-pass membrane receptor lacking selective antagonists, Z-TAC mediated complete receptor degradation and functional inhibition, demonstrating the ability of this platform to overcome the limitations of conventional pharmacological approaches. This study delineates a generalizable and scalable strategy for functional perturbation of the cell-surface proteome.

## Main

Cell-surface protein degraders have emerged as powerful tools for eliminating extracellular and membrane proteins^1,2^. Platforms such as LYTAC, AbTAC, KineTAC, EndoTag, and our previously reported TransTAC use bispecific molecules to recruit internalization receptors and drive endocytosis and lysosomal degradation of proteins of interest (POIs)^3–12^. As a result, they enable targeted removal of proteins that are difficult to modulate with conventional occupancy-based pharmacology, such as those with high endogenous ligand affinity and local concentrations, constitutive activity, or scaffolding functions^1,2,13–15^.

To date, most cell-surface degradation platforms require target-specific bispecific antibody engineering, requiring sequence information of the target binders and tailored design and optimization for each degrader, which curtails the scalability of these technologies^1,2^. Consistent with this, the range of targets explored has remained relatively narrow, mostly focusing on single-pass classical transmembrane proteins^8,16^. In addition, approaches for combinatorial protein degradation or for effective targeting and inhibition of multipass membrane proteins remain limited, constraining our ability to modulate the cell-surface protein networks^17^. As a result, widespread deployment of membrane protein degrader technologies has been constrained by the need for custom engineering, creating a barrier for laboratories without dedicated protein engineering expertise.

With decades of academic and industrial efforts, a vast repertoire of IgG antibodies has already been developed for research and clinical applications and are commercially available^18^. A strategy that could directly harness these existing antibodies for antigen modulation without requiring re-engineering into degrader formats would substantially broaden the range of degradable cell-surface targets. Such an approach would enable rapid exploration of diverse biological roles and therapeutic applications across various cell surface targets.

To develop a universal cell-surface protein degradation technology, we designed Z-TACs, which consist of a compact affinity-tuned TfR1-binding nanobody fused to an Fc-binding Z domain derived from *Staphylococcal* protein A^19,20^ (**Fig. 1a**). Z-TACs leverage protein A binding to the Fc region of IgG antibodies or Fc fusion ligands and harness transferrin receptor 1 (TfR1), a constitutively and rapidly internalizing effector widely expressed in many cell types^3,21,22^, to drive co-internalization at the cell surface and lysosomal degradation (**Fig. 1a**). Because TfR1 is broadly expressed across many cell and tissue types, this design enables degradation of targets in a wide range of endogenous cellular contexts. The IgG antibodies can originate from commercial sources, therapeutic antibodies, or in-house generated antibodies. The recombinant Z-TAC protein can be robustly expressed with high yield and purified both from mammalian cells (**Fig. 1b**) and *E. coli* (**Extended Data Fig. 1a**).

**Figure 1.**
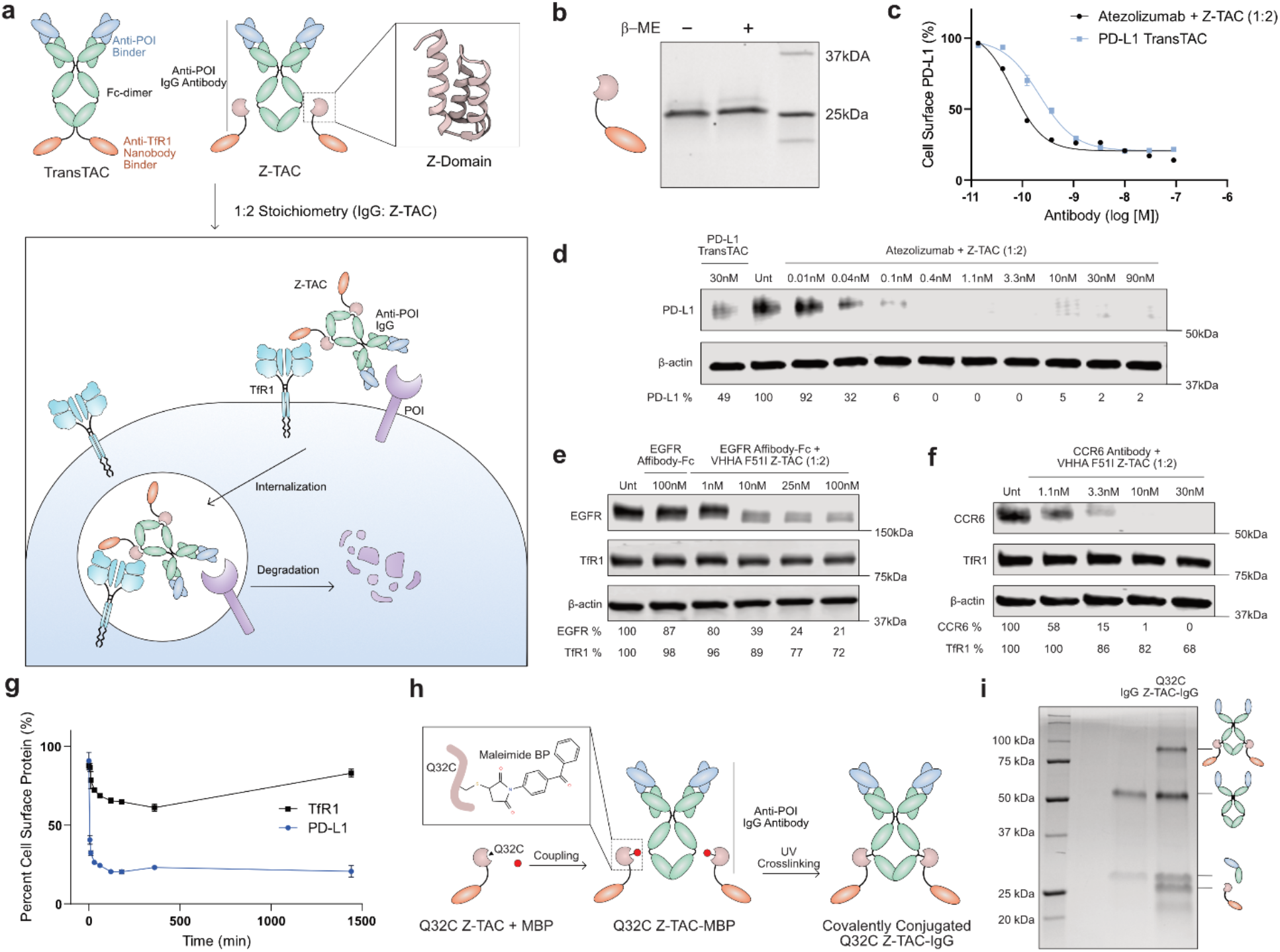
Z-TAC is a flexible platform for cell surface protein degradation using commercial, recombinant, and therapeutic IgG antibodies. **(a)** Schematic comparing TransTACs and Z-TACs. TransTACs are bispecific molecules that co-engage a POI antibody and TfR1 to induce lysosomal degradation. Z-TACs replace the engineered anti-POI arm with a compact Fc-binding Z-domain derived from Staphylococcal protein A, enabling receptor-mediated internalization and lysosomal degradation of the bound POI. **(b)** Purification of recombinant Z-TAC protein from Expi293 cells. SDS-PAGE analysis under non-reducing and reducing conditions shows the expected bands corresponding to monomeric Z-TAC (22.2 kDa). **(c)** Quantification of cell-surface PD-L1 levels by flow cytometry in MDA-MB-231 cells after 2 h treatment with indicated constructs and concentrations. Data represents means ± s.d. from n = 2 replicates. **(d)** Western blot analysis of PD-L1 degradation in MDA-MB-231 cells following treatment with atezolizumab + Z-TAC (1:2 stoichiometry) for 16 h compared with TransTAC and untreated controls. β-actin serves as a loading control. Remaining PD-L1 (% of untreated) is indicated below lanes. **(e)** Western blot analysis of EGFR degradation and TfR1 levels in PC9 cells following treatment with EGFR Affibody-Fc + VHHA F51I Z-TAC (1:2 stoichiometry) for 16 h compared with EGFR Affibody-Fc and untreated controls. β-actin serves as a loading control. Remaining EGFR (% of untreated) is indicated below lanes. **(f)** Western blot analysis of CCR6 degradation and TfR1 levels CCR6^+^ Jurkat cells following treatment with CCR6 antibody + VHHA F51I Z-TAC (1:2 stoichiometry) for 16 h compared with CCR6 antibody control. β-actin serves as a loading control. Remaining CCR6 (% of untreated) is indicated below lanes. **(g)** Time-course flow cytometry quantification of cell-surface PD-L1 and TfR1 following treatment with the Y96A Z-TAC variant and atezolizumab. Data represents means ± s.d. from n = 2 replicates. **(h)** Schematic illustrating a covalent conjugation strategy to generate Z-TAC-IgGs. **(i)** Reducing SDS-PAGE analysis of cetuximab and covalently conjugated Q32C Z-TAC-cetuximab.

We first used Z-TAC to internalize and degrade PD-L1, an immune checkpoint ligand with significant clinical interest^23^. The clinically approved anti-PD-L1 antibody atezolizumab was used as the IgG^24,25^. We quantified cell-surface PD-L1 levels in MDA-MB-231 cells following treatment with Z-TAC complexed with atezolizumab and compared its activity to that of a PD-L1 TransTAC^3^. Flow cytometry indicated dose-dependent depletion of surface PD-L1 (**Fig. 1c**). Z-TAC plus atezolizumab exhibited an IC_50_ of 66 pM, compared to 210 pM for the corresponding TransTAC, with both approaches achieving up to 80% depletion within 2 h (**Fig. 1c**). High levels of internalization were maintained across a range of PD-L1 atezolizumab:Z-TAC stoichiometries (**Extended Data Fig. 1b**). Consistent with surface loss of PD-L1, western blot analysis demonstrated complete PD-L1 degradation following 16 h of treatment with atezolizumab in combination with Z-TAC (**Fig. 1d**).

To establish the modularity of the Z-TAC platform and enable fine control over protein binding kinetics and degradation efficacy, we generated three Z-TAC variants spanning a range of TfR1 binding affinities^26^ and evaluated whether the technology is generalizable across different classes of cell-surface proteins. This included the parental VHHA Z-TAC (K_d_ < 1 nM), VHHA Y96A Z-TAC (K_d_ ∼ 3 nM), and VHHA F51I Z-TAC (K_d_ ∼ 124 nM) (**Extended Data Fig. 1c**). Using target-specific antibodies, we observed efficient degradation of multiple receptors in addition to PD-L1, including EGFR, a single-pass cancer-associated receptor tyrosine kinase (**Fig. 1e; Extended Data Fig. 2a–c**), and CCR6, a 7-span chemokine receptor implicated in immune dysregulations (**Fig. 1f; Extended Data Fig. 2d**). Notably, pre-incubation of the CCR6 antibody with Z-TAC did not measurably affect internalization efficacy, indicating rapid association of the Z-TAC with the IgG (**Extended Data Fig. 2f**). Across these targets and variants, Z-TAC treatment resulted in substantial depletion of receptor levels, demonstrating that the platform can accommodate structurally and functionally diverse membrane proteins.

While all affinity variants promoted target degradation, differences were observed in degradation potency and in the extent of TfR1 co-depletion. Across the targets examined, the intermediate-affinity VHHA Y96A construct provided the most favorable balance of robust degradation with limited TfR1 loss (**Fig. 2b; Extended Data Fig. 2b, d**). In contrast, the higher-affinity parental VHHA variant showed greater apparent potency but was associated with increased TfR1 depletion and a hook effect at higher concentrations (**Fig. 2d, Extended Data Fig. 2a, c**), a phenomenon also reported for small-molecule bispecific degraders^14^ and likely driven here by reduced TfR1 decoupling in endosomes. The lowest-affinity variant displayed reduced degradation potency and minimal impact on TfR1 levels (**Fig. 1e, f, Extended Data Fig. 1e**). These results indicate that TfR1 binding affinity influences both the efficiency of POI engagement and the extent to which POI trafficking can be decoupled from TfR1 recycling pathways. These Z-TAC variants constitute a toolbox that can be selectively deployed for different applications.

**Figure 2.**
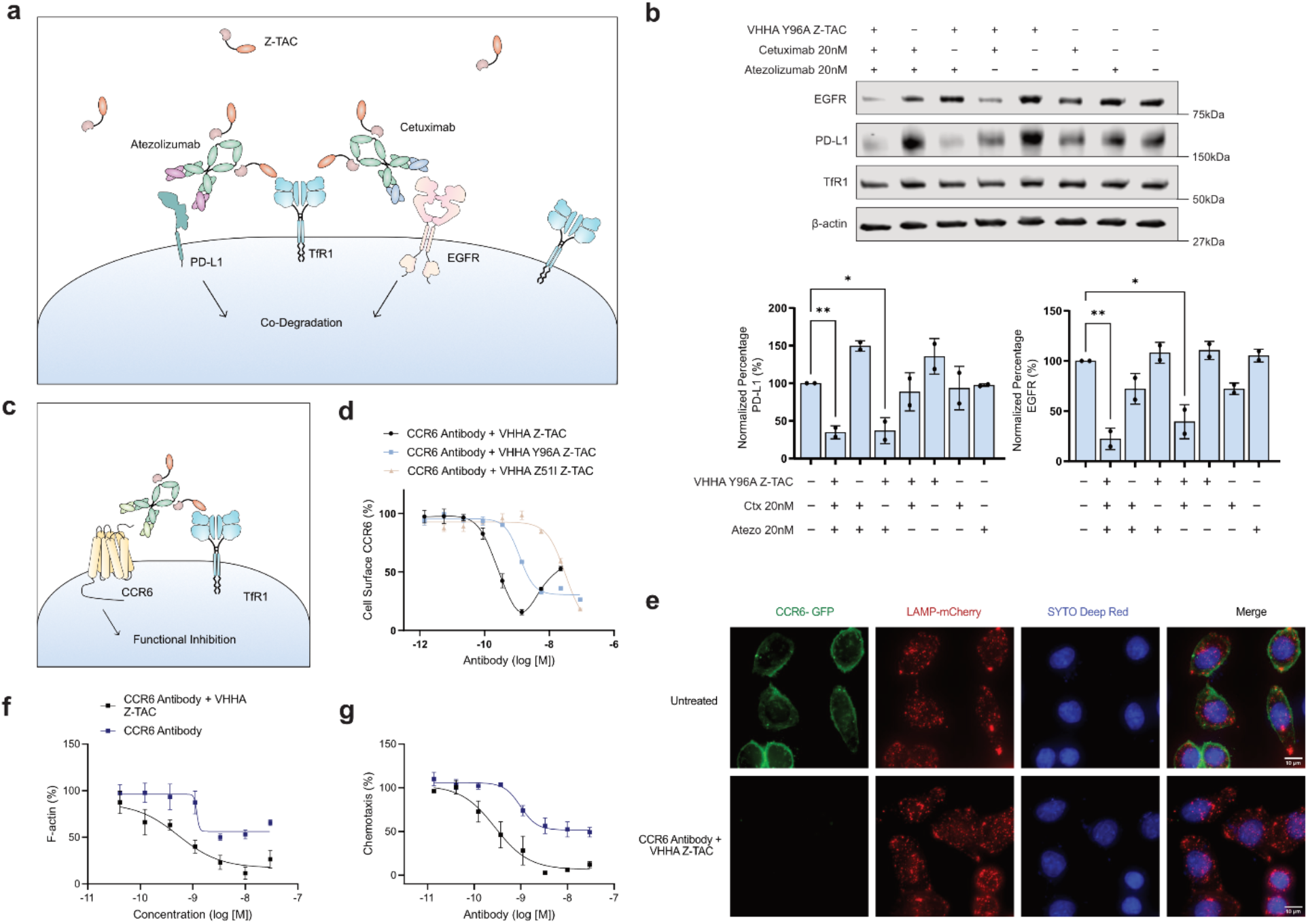
Z-TACs enable multiplex antibody-guided degradation and functional suppression of multi-pass membrane receptors. **(a)** Schematic illustrating that Z-TACs can be deployed to simultaneously co-degrade multiple targets when more than one IgG is present. **(b)** Western blot and quantifications showing Z-TAC mediated EGFR and PD-L1 co-degradation in MDA-MB-231 cells. Cells are treated with cetuximab (anti-EGFR), atezolizumab (anti-PD-L1), and VHHA F51I Z-TAC (1:2 stoichiometry) or controls for 16 h. β-actin serves as a loading control. **(c)** Schematic illustrating Z-TAC mediated functional inhibition of CCR6 via targeted degradation. **(b)** Flow cytometry dose-response quantification of cell-surface CCR6 levels in CCR6^+^ Jurkat cells after 2 h treatment with indicated constructs and concentrations. Data represents means ± s.d. from n = 2 replicates. **(e)** Microscopy images of HeLa cells expressing LAMP-mCherry and CCR6-GFP with 16 h treatment of 10 nM anti-CCR6 antibody and 20nM VHHA Z-TAC **(f)** Flow cytometry quantification of total F-actin in CCR6^+^ Jurkat cells with fixation and phalloidin staining. Cells were treated with varying the indicated constructs for 2 h, followed by 1 min stimulation with 50 nM CCL20. Data represents means ± s.d. from n = 3 replicates. **(g)** Chemotaxis of CCR6^+^ luciferase^+^ Jurkat cells measured by Transwell migration assay after 2 h treatment & 2 h migration with CCL20 as the chemoattractant. Data represents means ± s.d. from n = 3 replicates.

We next performed time-course flow cytometry to define the kinetics of Z-TAC-mediated internalization in MDA-MB-231 cells. Cells were treated with the anti-PD-L1 antibody atezolizumab together with the VHHA Y96A Z-TAC variant. After treatment, cell-surface PD-L1 levels decreased rapidly, with greater than 70% depletion observed within only 5 minutes and sustained suppression over 24 h (**Fig. 1g**). In contrast, TfR1 exhibited only a transient reduction followed by recovery over 24 h, consistent with receptor recycling and supporting selective depletion of the target protein without significantly altering TfR1 homeostasis. These results show that Z-TAC-mediated POI degradation is rapid and sustained.

Although the mix-and-use format of Z-TACs enables highly modular targeting using existing antibodies, certain applications may benefit from a covalently conjugated antibody–Z-TAC format, which can enhance complex stability and ensure defined stoichiometry, particularly during prolonged incubation or *in vivo* where noncovalent interactions may dissociate or endogenous IgG may compete for Z-TAC binding. To enable this, we utilized a covalent conjugation strategy to generate stable Z-TAC-IgG complexes^27^. In this approach, a cysteine-engineered Z domain variant (Q32C Z-TAC) was first modified through thiol-maleimide coupling with a maleimide-benzophenone (MBP) crosslinker to generate Q32C Z-TAC-MBP. Upon binding of the Z domain to the Fc region of an IgG antibody, UV irradiation activates the benzophenone moiety, which forms a covalent crosslink between the Z-TAC and the antibody Fc (**Fig. 1h**). Formation of the covalent conjugate was confirmed by reducing SDS-PAGE, which showed a molecular weight shift of the IgG heavy chain following conjugation (**Fig. 1i**).

The modular architecture of Z-TACs also creates opportunities for combinatorial targeting of cell-surface proteins (**Fig. 2a**). Many signaling pathways are governed by coordinated receptor networks rather than single targets, making simultaneous modulation of multiple receptors an attractive strategy for both biological discovery and therapeutic development^28,29^. To evaluate this capability, we treated cells with a combination of cetuximab, an anti-EGFR antibody, and atezolizumab, an anti-PD-L1 antibody, in the presence of VHHA Y96A Z-TAC^30^. Co-treatment resulted in efficient degradation of both receptors without evidence of interference between targeting events, demonstrating that Z-TAC-mediated trafficking through TfR1 can accommodate multiple antibody-POI complexes (**Fig. 2b**). These results validate the feasibility of combinatorial degradation using the Z-TAC platform and highlight its potential for multiplexed modulation of cell-surface signaling networks.

One application of protein degraders is functional interrogation and therapeutic targeting of receptors for which effective or selective antagonists are unavailable. CCR6 is a chemokine receptor within the G-protein coupled receptor (GPCR) family that regulates immune cell trafficking and inflammatory signaling. It represents a compelling drug target across multiple autoimmune and inflammatory diseases, yet selective pharmacological inhibitors remain limited^31,32^. Existing small-molecule inhibitors lack specificity and exhibit cross-reactivity with other chemokine receptors^33,34^. Antibody-based antagonists achieve only partial pathway inhibition, typically reaching a maximum of ∼50-60%, because the endogenous ligand CCL20 binds CCR6 with high affinity and engages multiple receptor epitopes^35^, making competitive blockade inherently challenging^36^.

We therefore reasoned that Z-TAC-mediated degradation of CCR6 could provide a more effective means of suppressing receptor signaling by eliminating the receptor from the cell surface rather than competing for ligand binding (**Fig. 2c**). We first evaluated CCR6 internalization induced by different Z-TAC variants in combination with an anti-CCR6 antagonist antibody. Flow cytometry revealed robust, dose-dependent receptor internalization across variants. The VHHA Y96A Z-TAC exhibited potent activity with an IC_50_ of 1.2 nM and the VHHA Z51I Z-TAC displayed an IC_50_ of 31 nM (**Fig. 2d**). To visualize receptor loss in living cells, we performed fluorescence imaging of CCR6-GFP following treatment with antibody Z-TAC complexes. After 16 h, cells treated with VHHA Z-TAC showed a pronounced reduction in CCR6 signal compared with controls, consistent with receptor degradation (**Fig. 2e**).

Activation of CCR6 initiates downstream F-actin-mediated cytoskeletal remodeling and cell migration. In F-actin polymerization assays, treatment with antibody–VHHA Z-TAC complexes for 2 h resulted in potent suppression of CCL20-induced actin polymerization, reaching up to 85% inhibition with an IC_50_ of 0.5 nM, substantially exceeded that achieved with the antibody alone (**Fig. 2f**)^37^. Moreover, in Transwell chemotaxis assays, Z-TAC treatment produced near-complete suppression of CCR6-mediated Jurkat cell migration, achieving up to 95% inhibition with an IC_50_ value of 0.3 nM, whereas treatment with the antibody alone resulted in up to ∼50% inhibition (**Fig. 2g**). These findings demonstrate that receptor degradation can enhance functional antagonism of GPCR signaling and provide an effective strategy for modulating receptor pathways that are challenging to control using conventional occupancy-based approaches.

## Discussion

Cell-surface proteins represent a large and biologically important class of therapeutic targets, yet many remain difficult to modulate pharmacologically. Z-TAC offers a modular strategy for converting existing binders into degraders. Because the platform does not require target-specific engineering, it enables rapid repurposing of the extensive repertoire of antibodies developed over decades of academic and industrial research to generate functional modulators of diverse cell-surface targets. This capability is particularly valuable for proteins for which selective and potent antagonists are unavailable, a challenge that applies to a substantial fraction of membrane proteins, such as including many GPCRs, ion channels, transporters, and receptors with constitutive activity^1,2^. Compared with genetic perturbation strategies, Z-TAC enables post-translational control of protein function and allows interrogation of cellular responses to receptor perturbation within the endogenous cellular context.

Z-TACs are straightforward to express, purify, and deploy. They can be produced in *E. coli* or mammalian Expi293 cells with high yield, and cellular assays require only simple mixing with an antibody, making the platform accessible to a wide range of laboratories. For applications in more complex systems, we implemented a covalent conjugation strategy that converts the noncovalent complex into a stable conjugate, preventing dissociation or unintended binding to other IgG molecules. The plug-and-play design also enables simple implementation of combinatorial degradation, allowing simultaneous modulation of multiple cell-surface targets to interrogate cellular networks.

We evaluated Z-TAC across three representative cell-surface targets spanning distinct protein classes, including PD-L1, an immune checkpoint ligand; EGFR, a receptor tyrosine kinase; and CCR6, a chemokine receptor GPCR. Despite differences in receptor structure, endocytic properties, and signaling mechanisms, Z-TAC induced efficient degradation across all targets tested to date without the need for target-specific engineering or optimization. These findings support the broad generalizability of the platform and highlight the potential of Z-TAC to enable systematic functional interrogation of cell-surface protein biology.

## Methods

### Plasmid construction

Plasmids were constructed using standard Gibson cloning methods. For binders, DNA fragments of TfR1 VHHs, Z domain, anti-EGFR affibody, and CCR6 VH/VL domains, were synthesized by Integrated DNA Technologies (IDT). Anti-human EGFR plasmid was obtained from Addgene. Binder sequences were subcloned into a pFuse-hIgG1 vector (InvivoGen) for mammalian expression or a pBR322 vector for E Coli expression. Eleven- or 12-amino acid linkers were inserted between binders. An N-terminal 8xHis-tag was added to some constructs for purification.

### Cell lines and culture

Lenti-X 293T and MDA-MB-231 cells were cultured in DMEM (ThermoFisher Scientific) with 10% Fetal Bovine Serum (FBS) and 1% penicillin/streptomycin, incubated at 37°C with 5% CO2. Jurkat and PC9 cells were cultured in RPMI-1640 (ThermoFisher Scientific) with 10% FBS and 1% penicillin/streptomycin, incubated at 37°C with 5% CO2. All other Expi293 cells were cultured in Expi293 Expression Medium (ThermoFisher Scientific), incubated with gentle shaking at 37°C with 8% CO2.

### Protein expression

Antibodies and Z-TAC constructs were expressed in Expi293 cells (ThermoFisher Scientific) using FectoPRO (ThermoFisher Scientific) according to the manufacturer’s protocol. Antibodies were purified by Protein A affinity chromatography and buffer exchanged into PBS. Z-TAC constructs were also expressed in *E. coli* C43 (DE3) cells (Sigma-Aldrich). Plasmids were transformed by heat shock and cultured in 2XYT medium containing ampicillin (100 µg mL^-1^) and chloramphenicol (12.5 µg mL^-1^). Expression cultures were induced at OD600 0.6–0.8 with IPTG and incubated overnight at 18 °C with shaking. Cells were harvested by centrifugation, lysed using B-PER supplemented with DNase I, and clarified by centrifugation. Z-TACs were purified by Ni-NTA affinity chromatography and buffer exchanged into PBS. Proteins were kept at 4°C for up to 1-2 weeks or flash frozen for storage at -80°C. The purity and integrity of proteins were confirmed by SDS-PAGE electrophoresis.

### Lentiviral packaging and engineered cell generation

For lentivirus production, Lenti-X 293T cells were grown to 70–80% confluency before the medium was replaced with fresh DMEM containing 15% FBS. Cells were transfected with the carrier plasmid and lentiviral helper plasmids (psPax2 and pMD2.G) using a standard Lipofectamine3000 protocol (Thermo Fisher Scientific). The following day, the medium was replaced with fresh DMEM containing 20% FBS. 48-60 h post-transfection, the supernatant containing lentivirus was collected and centrifuged at 15,000 g for 1 min to remove cell debris. If not used right away, lentivirus was flash frozen with dry ice prior to storage at -80°C.

Jurkat cells were plated at a density of 0.75 × 10^6^ cells/mL and were then infected with doxycycline-inducible FLAG-CCR6 lentivirus diluted 1:7 with 8 ug/mL polybrene via spinfection spinning at 890g at 30°C for 2 h. The medium was changed the next day with 20% FBS RPMI-1640, and cells were further cultured and expanded before selecting for CCR6+ Jurkat cells with the addition of 2 ug/mL puromycin.

### General setups for internalization or degradation experiments

Suspension cells were seeded at 0.6 × 10^6^ cells per millilitre in complete growth medium and Z-TACs with IgGs or controls were added immediately to the cells at specified concentrations; adherent cells were seeded and grown for 6–12 h to reach roughly 90% confluency before TransTACs or controls were added to the cells. For all assays, cells were incubated with Z-TACs with IgGs or controls for 2–24 h before flow cytometry, western blotting or microscopy experiments, unless otherwise noted.

### Western blotting

Cells were lysed with 1x RIPA buffer and 1x protease inhibitor (ThermoFisher) at 4 °C for 20 min. For CCR6, cells were lysed in lysis buffer consisting of 1% digitonin, 150 mM NaCl, and 100 mM Tris pH 7.5 and 1x protease inhibitor at 4 °C for 40 min. Lysates were centrifuged at 16,000*g* for 10 min at 4 °C. Supernatants were mixed with 4× Laemmli Sample Buffer (Bio-Rad) and 2-mercaptoethanol and heated at 95 °C for 5 min. For CCR6, 2-mercaptoethanol was not added and samples were not heated. Lysates were not boiled and were directly loaded on SDS–PAGE gels (Bio-Rad). Proteins were transferred to polyvinylidene difluoride membranes using a Trans-Blot Turbo Western Blotting Transfer System (Bio-Rad), followed by standard Western Blot protocols for washing and primary/secondary antibody staining. Membranes were imaged with a LICOR imager and Image Studio software (v.5.2.5). Band intensities were quantified using ImageJ (v.2.16). Band intensities were quantified using ImageJ (v.2.16). Antibodies used included anti-PD-L1 (Cell Signaling Technology, 13684T, 1:1,000), anti-EGFR (Biolegend, 933901, 1:2500), anti-β-actin (Cell Signaling Technology, 4970S, 1:1,000), anti-TfR1 (CD71) (Cell Signaling Technology, 13113S, 1:1,000), anti-CCR6 (ThermoFisher, 14-1969-82, 1:300), anti-TfR1 (ThermoFisher, 13-6800, 1:1000), IRDye 800CW Goat anti-mouse IgG (LICOR, 926-32210, 1:10,000), IRDye 680RD goat anti-rabbit IgG (LICOR, 926-68071, 1:10,000).

### Flow cytometry

Harvested cells were centrifuged at 300 g for 5 min. Pellets were washed 1x with FACS buffer (ice-cold PBS + 3% BSA). Cells were incubated with primary antibody in FACS buffer for 20-30 min on ice. Cells were washed 3 times and resuspended in FACS buffer for flow analysis. Flow cytometry was performed using a CytoFLEX cytometer (Beckman Coulter, V0-B2-R2)) and CytExpert software (v2.3.1.22). FACS was performed using a CytoFLEX SRT cell sorter (Beckman Coulter, V0-B2- Y5-R3) and CytExpert SRT software (v.1.1.0.10007). Data was analyzed with FlowJo (v10.8.0).

Primary antibodies include: Alexa Fluor 647 anti-FLAG Tag (M1) (1:500, purified from hybridoma supernatant and conjugated in-house), FITC anti-human CD71 (Biolegend, 334104, 1:200), and Alexa Fluor 647 anti-PD-L1 (Cell Signaling Technologies, 41726S,1:100)

### Maleimide benzophenone modification

Z-TAC proteins were modified with 4-(N-maleimido)benzophenone (MBP) (MedChemExpress) to enable photo-crosslinking to IgG antibodies. Purified Z-TAC was first concentrated to 2.5 mg mL^-1^ in PBS and reduced in the presence of tris(2-carboxyethyl)phosphine (TCEP) and EDTA (5 mM) for 30 min at 37 °C. MBP was dissolved in DMSO to 18mM and added to the reduced protein at a final concentration of 500 µM, followed by incubation in the dark at 37 °C for 45 min to allow thiol–maleimide coupling. The reaction was quenched with excess cysteine and the modified protein (Z-TAC–MBP) was buffer exchanged into PBS using PD MiniTrap™ G-10 columns (Cytiva).

### Photoconjugation of IgG antibodies

IgG antibodies were mixed with Z-TAC–MBP at a molar excess of Z-TAC–MBP and incubated for 30 min at room temperature in the dark to allow Fc binding. Samples were then exposed to 365 nm UV light for 2 h on ice using a UV crosslinker (Analytik Jena) to activate the benzophenone moiety and induce covalent crosslinking to the antibody Fc region. Following UV crosslinking, samples were incubated with excess Protein A beads to capture unconjugated IgGs. The unbound fraction was collected and analyzed by SDS-PAGE electrophoresis.

### F-Actin polymerization

Doxycycline-inducible CCR6+ Jurkat cells were treated with 25 ng/ul doxycycline. After 24 h, 2.5 × 10^5^ cells were seeded in a 96-well flat bottom plate and were incubated with GTACs for 2 h. 10 nM of GTACs were used for experiments with only one dose. Cells were then resuspended in a V-bottom plate with 50 nM CCL20 followed by staining and fixation with phalloidin staining buffer after the indicated time. Phalloidin staining buffer consists of 1% BSA, 0.2% Triton X-100, 4% paraformaldehyde, and 1 U of Flash Phalloidin Green 488 (BioLegend). Cells were stimulated for 1 min for experiments with only one time point. Cells were washed 3x and resuspended in cold 0.1% Triton X-100 in PBS for flow cytometry analysis.

### Chemotaxis assay

Doxycycline-inducible CCR6+ luciferase+ Jurkat cells were treated with 25 ng/ul doxycycline. After 24 h, 4 × 10^5^ cells were seeded in a 96-well flat bottom plate and were incubated with GTACs for 2 h in 100 ul of serum-free RPMI-1640. Each 100 ul cell solution was then transferred to Hanging Cell Culture inserts with 5 µm pores (Millipore Sigma). These inserts were placed within a 24-well plate which contained 600 ul of 100 nM CCL20, and cells were allowed to incubate for 2 h. Then, migrated cells were collected from the outer chambers of the 24-well plate and were transferred to a white 96-well flat-bottom plate. 150 µg/ml D-luciferin (GoldBio) was added to the cells to allow for quantification via luminescent readout with a CLARIOstar microplate reader (BMG Labtech).

### Live-cell imaging

4 × 10^4^ HeLa cells were plated in poly-D-lysine-coated 24-well glass bottom plates 2 days prior to imaging. Cells were treated for 16 h, stained with SYTO™ Deep Red nucleic acid stain for live cells for 30 min (ThermoFisher Scientific), and washed three times with PBS before imaging. Cells were imaged using a LEICA DMi8 fluorescence microscope (Leica Application Suite X v3.7.5.24914). Images were processed and analyzed using ImageJ (v.2.16).

### Data and statistical analysis

All graphing and statistical analyses were performed in GraphPad Prism (v10.4.2) or Microsoft Excel (v16.98). Unpaired two-tailed t-tests were used for statistical analyses unless otherwise noted. A two-tailed P-value was used to determine statistical significance for all analyses. P < 0.05 was considered statistically significant. ns: P>0.05, *: P<= 0.05, **: P<=0.01, ***: P <=0.001, ****: P<=0.0001. Figures were created with BioRender.com and Affinity Designer (v2.6.3).

## Acknowledgment

We thank Rebecca Metivier for assistance with mass spectrometry analysis and members of the Flynn and Zhou lab for helpful discussion. Fig. 1a was created with BioRender.com.

---

**Extended Data Fig. 1.**
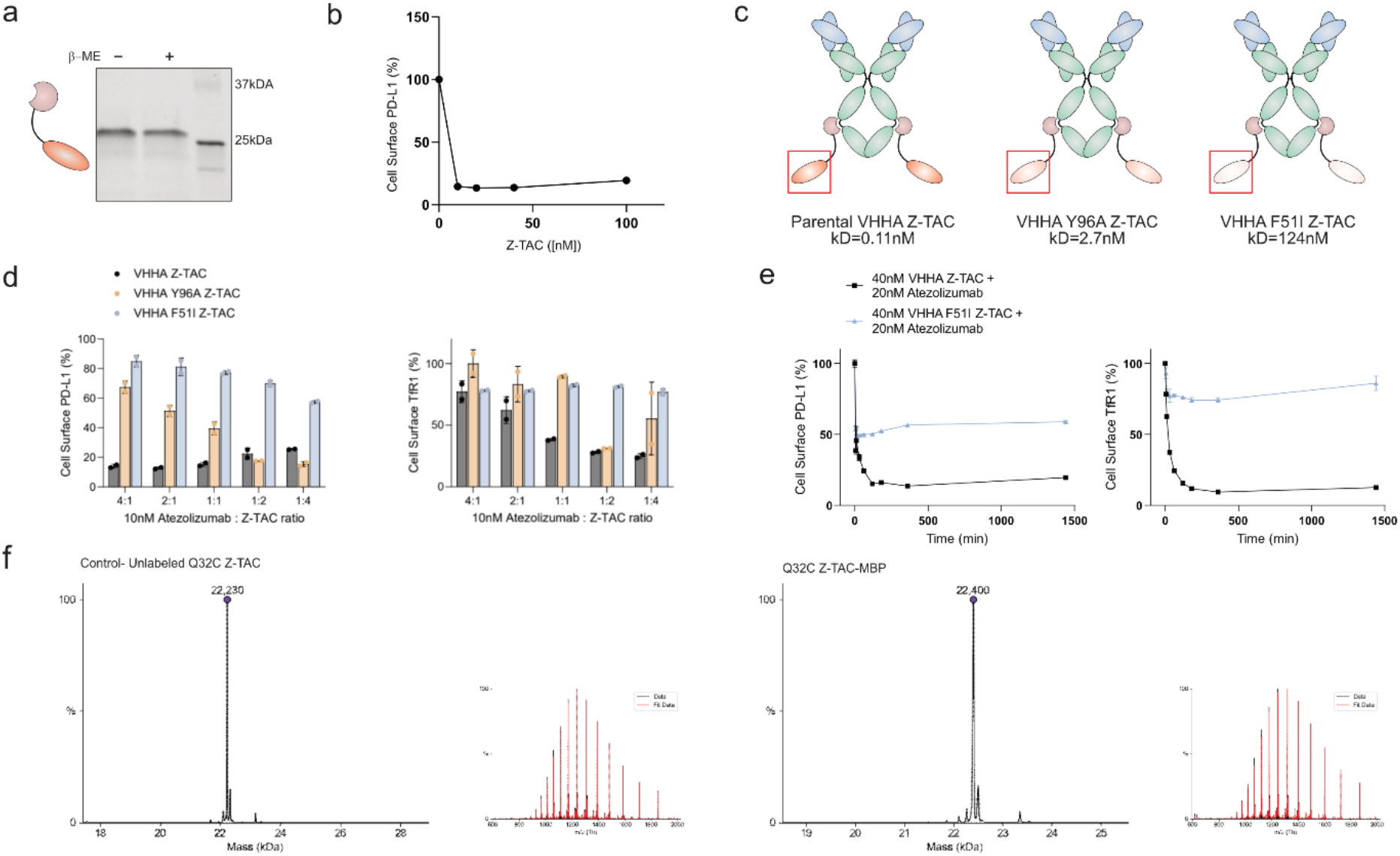
Variant Z-TAC toolbox with different TfR1 affinities. **(a)** Purification of recombinant Z-TAC protein from E. coli. SDS-PAGE analysis under non-reducing and reducing conditions shows the expected bands corresponding to monomeric Z-TAC (∼23 kDa). **(b)** Quantification of cell-surface PD-L1 levels by flow cytometry in MDA-MB-231 cells after 2 h treatment with different Z-TAC concentrations in the presence of 20nM atezolizumab by flow cytometry. Data represents means ± s.d. from n = 2 replicates. **(c)** Schematic of a Z-TAC “toolbox” consisting of variants with different affinity to TfR1. **(d)** Quantification of cell-surface PD-L1 (left) and TfR1 (right) levels by flow cytometry in MDA-MB-231 cells after 2 h treatment with varying atezolizumab: Z-TAC ratio for the indicated Z-TAC variants. Data represents means ± s.d. from n = 2 replicates. **(e)** Time-course quantification of cell-surface PD-L1 (left) and TfR1 (right) levels by flow cytometry in MDA-MB-231 cells comparing 40nM parental VHHA Z-TAC and the lower-affinity VHHA F51I Z-TAC in the presence of 20nM atezolizumab. Data represents means ± s.d. from n = 2 replicates. **(f)** Intact mass spectrometry confirming complete modification of Q32C Z-TAC with maleimide-benzophenone.

**Extended Data Fig. 2.**
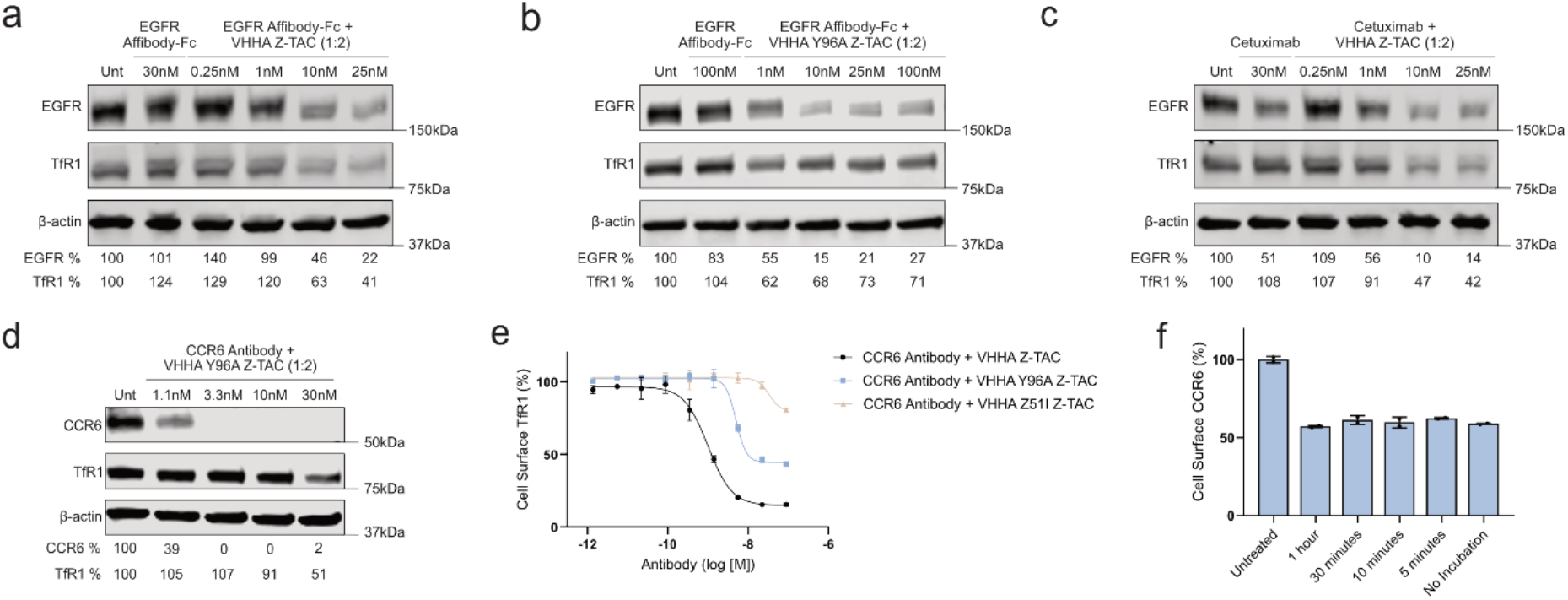
EGFR and CCR6 internalization and degradation by Z-TAC across affinity variants. **(a)** Western blot analysis of EGFR degradation and TfR1 levels in PC9 cells following treatment with EGFR Affibody-Fc + VHHA Z-TAC (1:2 stoichiometry) for 16 h compared with EGFR Affibody-Fc and untreated controls. β-actin serves as a loading control. Remaining EGFR (% of untreated) is indicated below lanes. **(b)** Western blot analysis of EGFR degradation and TfR1 levels in PC9 cells following treatment with EGFR Affibody-Fc + VHHA Y96A Z-TAC (1:2 stoichiometry) for 16 h compared with EGFR Affibody-Fc and untreated controls. β-actin serves as a loading control. Remaining EGFR (% of untreated) is indicated below lanes. **(c)** Western blot analysis of EGFR degradation and TfR1 levels in PC9 cells following treatment with cetuximab + VHHA Z-TAC (1:2 stoichiometry) for 16 h compared with cetuximab and untreated controls. β-actin serves as a loading control. Remaining EGFR (% of untreated) is indicated below lanes. **(d)** Western blot analysis of CCR6 degradation and TfR1 levels CCR6^+^ Jurkat cells following treatment with CCR6 antibody + VHHA Y96A Z-TAC (1:2 stoichiometry) for 16 h compared with CCR6 antibody control. β-actin serves as a loading control. Remaining CCR6 (% of untreated) is indicated below lanes. **(e)** Flow cytometry dose-response quantification of cell-surface TfR1 levels in CCR6^+^ Jurkat cells after 2 h treatment with indicated constructs and concentrations. Data represents means ± s.d. from n = 2 replicates. **(f)** Bar graph representation of cell-surface CCR6 across the indicated Z-TAC and antibody pre-incubation times, measured by flow cytometry following treatment with CCR6 antibody plus VHHA Z-TAC (1:2 stoichiometry) for 2 h. Data represent means ± s.d. from n = 2 replicates.

